# Differential effects of antibiotic treatment with piperacillin/tazobactam or ceftriaxone on the murine gut microbiota

**DOI:** 10.1101/2020.05.28.122473

**Authors:** Carola Venturini, Bethany Bowring, Alicia Fajardo-Lubian, Carol Devine, Jonathan Iredell

**Affiliations:** Centre for Infectious Diseases and Microbiology, Westmead Institute for Medical Research (WIMR), Sydney, Australia; School of Medicine, Sydney Medical School, University of Sydney, Australia; Westmead Hospital, Western Sydney Local Health District (WSLHD), Sydney, Australia

## Abstract

Effective antimicrobial stewardship requires a better understanding of the impact of different antibiotics on the gut microflora. Studies in humans are confounded by large inter-individual variability and difficulty in identifying control cohorts. However, controlled murine models can provide valuable information. We examined the impact of a penicillin-like antibiotic (piperacillin/tazobactam, TZP) or a third-generation cephalosporin (ceftriaxone, CRO) on the murine gut microbiota. We analyzed gut microbiome composition by 16S-rRNA amplicon sequencing and effects on the Enterobacteriaceae by qPCR and standard microbiology. Colonization resistance to multidrug resistant *Escherichia coli* ST131 and *Klebsiella pneumoniae* ST258 was also tested. Changes in microbiome composition and a significant (p<0.001) decrease in diversity occurred in all treated mice, but were more marked and longer lasting after CRO exposure with a persistent rise in Proteobacteria levels. Increases in the Enterobacteriaceae occurred in all antibiotic treated mice, but were transient and associated with direct antibiotic pressure. Co-habitation of treated and untreated mice attenuated the detrimental effect of antibiotics on treated animals, but also caused disturbance in untreated co-habitants. At the height of dysbiosis after antibiotic termination, the murine gut was highly susceptible to colonization with both multidrug resistant pathogens. The administration of a third-generation cephalosporin caused a significantly prolonged dysbiosis in the murine gut microflora, when compared to a penicillin/β-lactam inhibitor combination with comparable activity against medically important virulent bacteria. At the height of dysbiosis, both antibiotic treatments equally led to microbial imbalance associated with loss of resistance to gut colonization by antibiotic-resistant pathogens.

## Introduction

The current global health crisis in antibiotic resistance (AR) requires immediate action,^1,2^ and implementation of effective antibiotic stewardship strategies has become a major goal for health agencies worldwide. Optimal policy choices are ideally founded in a deep understanding of both the epidemiology and mechanisms of antimicrobial resistance, and the impact of different antibiotics on patient health. Exposure to antibiotics not only promotes the amplification and transfer of antibiotic resistance genes, but also induces a dysbiosis that facilitates colonization by opportunistic pathogens.^3–6^ The nature and duration of this dysbiosis have been shown to vary depending on the type of antibiotic used, and antibiotics with similar activity against principal human pathogens may have substantially different impacts on the intestinal microbiota of the gut.^5–9^

Proteobacterial blooms in the gut microflora, with increasing prevalence of Gammaproteobacteria (e.g. Enterobacteriaceae), have been associated with antibiotic treatment.^7,10,11^ However, the extent or duration of these blooms from different antibiotic treatments have not yet been fully investigated. The intestine is a particularly fertile environment for genetic exchange and the Enterobacteriaceae play a key role in AR dissemination.^4,11^ Antibiotic use provides direct selective pressure for AR spread through horizontal gene transfer mechanisms and stimulation of the SOS response and recombination,^12^ and plasmid transfer may be promoted in enterobacterial blooms.^4,13^ Both antibiotic resistant species and opportunistic pathogens thrive in a dysbiotic gut, and prolonged or repeated antibiotic therapy promotes enteric infections.^9,11,12,14,15^ *Escherichia coli* and *Klebsiella pneumoniae* are among the most important multidrug resistant (MDR) enteric opportunistic pathogens that may amplify during dysbiosis and cause life-threatening infections, particularly in high risk clinical settings (e.g. critical care units).^16,17^

Studies of microbiome dynamics in humans are confounded by large inter-individual variability, sampling size and difficulty in identifying control cohorts.^18,19^ Murine models of gut dysbiosis have been criticized for not being directly informative of the specific changes in human gut microbiome composition.^7,20^ Despite this, controlled experiments in these animal models provide valuable data on microbiome-wide effects that may be masked in human studies by inter-individual variability.^7,9,21,22^

Here, we examined the gut microbiota of mice treated with two antibiotics routinely used in emergency empiric antibiotic therapy: a penicillin-like antibiotic and β-lactamase inhibitor combination, piperacillin/tazobactam (Tazocin), and a third-generation cephalosporin, ceftriaxone. These antibiotics have comparable spectrum of activity but belong to different classes, and the third-generation cephalosporins (though not the penicillin) are notoriously associated with increased AR and pathogen colonization.^5,7,8,15,23–25^ We therefore also tested colonization resistance in the dysbiotic gut towards two MDR extended spectrum β-lactamase producing (ESBL) opportunistic pathogens, *E. coli* ST131 and *K. pneumoniae* ST258. We found that treatment with CRO was associated with more severe and prolonged dysbiosis, but that both antibiotics equally promoted the establishment of antibiotic-resistant pathogens.

## Materials and Methods

### Ethics

All animal experiments were approved by the appointed Animal Ethics Committee (ARA 4205.06.13; Western Sydney Local Health District, NSW Government), and were conducted in the Biosafety Animal Facility at the Westmead Institute for Medical Research (Sydney, Australia) complying with the national standards and guidelines for animal experimentation (*Australian code for the care and use of animals for scientific purposes* (2013), National Health and Medical Research Council (NHMRC), Australian Government).

### Experimental setup

The experimental workflow for this study is summarized in Figure S1. Faeces were considered as a proxy of intestinal microflora^21,26,27^ and, in all experiments, bacterial load was calculated as colony forming units (CFU) per gram of stool (CFU/g) (limit of detection 10_2_ CFU/g). In all instances, faecal pellets were weighed and homogenized in 0.9% saline by shaking (~15 min) with 6 mm glass beads (Benchmark Scientific, Sayreville, NJ, USA).

### Antibiotic treatment and microbiology

Female Balb-C mice (n=3 per group) were injected subcutaneously with either saline (group: ‘saline’), co-formulated piperacillin/tazobactam (Tazocin 6 mg / 750 ng; group: ‘TZP’) or ceftriaxone (2 mg/day; group: ‘CRO’) once a day for five consecutive days (day 1 = 24 h from first antibiotic injection) in daily weight-adjusted doses equivalent to those recommended for adult human patients.^28,29^ Recovery after antibiotic treatment was monitored for four weeks (day 9 to 32). Faecal pellets were collected every day during antibiotic therapy (day 1 to 5), and on selected days in recovery for microbiome analysis (Figure S1). We also exploited the coprophagic lifestyle of rodents to determine the effect of faecal transplantation on the recovery of dysbiotic mice by including a cohort of mice where one mouse treated with either antibiotic was co-housed with two untreated mice (groups: ‘co-TZP’ and ‘co-CRO’). An experimental control group (group: ‘none’, no treatment) was also included.

In order to exclude an effect from residual antimicrobial activity, we also measured antibiotic levels in homogenized faeces by modification of the CDS disc diffusion method, placing 9 mm paper disks (Schleicher & Schuell, Dassel, Germany) soaked in antimicrobials on lawns of sensitive bacteria (*E. coli* ATCC 25922) on Mueller-Hinton agar.^30^ The limit of detection was maximized by concentrating faecal resuspensions by vacuum centrifugation. The zone of clearing was compared to known concentrations of TZP and CRO similarly applied to the paper discs. Throughout the study, the total murine microbial flora was monitored by standard microbiology (growth on selective media): ChromAgar^™^ supplemented with vancomycin (20 μg/ml; Van_20_) for detection of total enterobacteria (no enterococci), and with Van_20_ plus cefotaxime (8 μg/ml; CTX_8_) for ESBL enterobacteria (Figure S2). Commensal *E. coli* and ESBL *E. coli* ST131 were identified on MacConkey agar (Becton Dickinson, Sparks, MD, USA) without and with CTX8, respectively (Figure S2). MacConkey agar supplemented with inositol (10 mM) and carbenicillin (100 μg/ml) was used for detection of *K. pneumoniae* ST258.^31^ Representative colonies with different morphology were typed using the Bruker MALDI Biotyper system (Bruker, Massachusetts, USA).

### Colonization resistance

Two MDR pathogens, common opportunistic colonizers of the gut, were selected for colonization experiments. ESBL *E. coli* ST131 [JIE3430 with *bla*_CTX-M-15_; MIC CRO >16 mg/L and MIC TZP ≤4/4 mg/L]^32^ and carbapenemase resistant *K. pneumoniae* ST258 [JIE2709; MIC CRO >16 mg/L and MIC TZP >64/4 mg/L]^33^ were routinely cultured in lysogeny broth (LB) at 37°C with shaking. Mice (n=3 per group) treated with either CRO or TZP (as above) were colonized orally on day 9 (recovery) with either *E. coli* or *K. pneumoniae* (10^9^ CFU/mL in 20% sucrose solution) as previously.^34^ Two control groups (n=3 each), that had not received any antibiotic treatment, were also colonized with either *E. coli* or *K. pneumoniae*. Water intake was monitored to ensure comparable amounts were consumed in each experimental group. Bacterial load was assessed for five weeks after first antibiotic administration by growth on selective MacConkey agar supplemented with cefotaxime (8 μg/mL; CTX8) only for *E. coli*, and with CTX8 plus 10 mM inositol plus 100 μg/ml carbenicillin for *K. pneumoniae*^31^ (Figure S2). To exclude the presence of other ESBL *E. coli* and *K. pneumoniae* in the murine gut, dilutions of faeces for each treatment group were plated on these selective media prior to colonization (and throughout the experiment for controls). To further confirm the identity of the *E. coli* and *K. pneumoniae* grown from faeces, we tested random colonies (n=8 each) by PCR (Table 1). Faecal pellets from day 1 (pre-antibiotic) and day 30 (endpoint, recovery week 4) were also plated on selective agar and total gut Gram-negative aerobic flora (CFU/g) quantified by standard microbiology techniques as above.

**Table 1.**
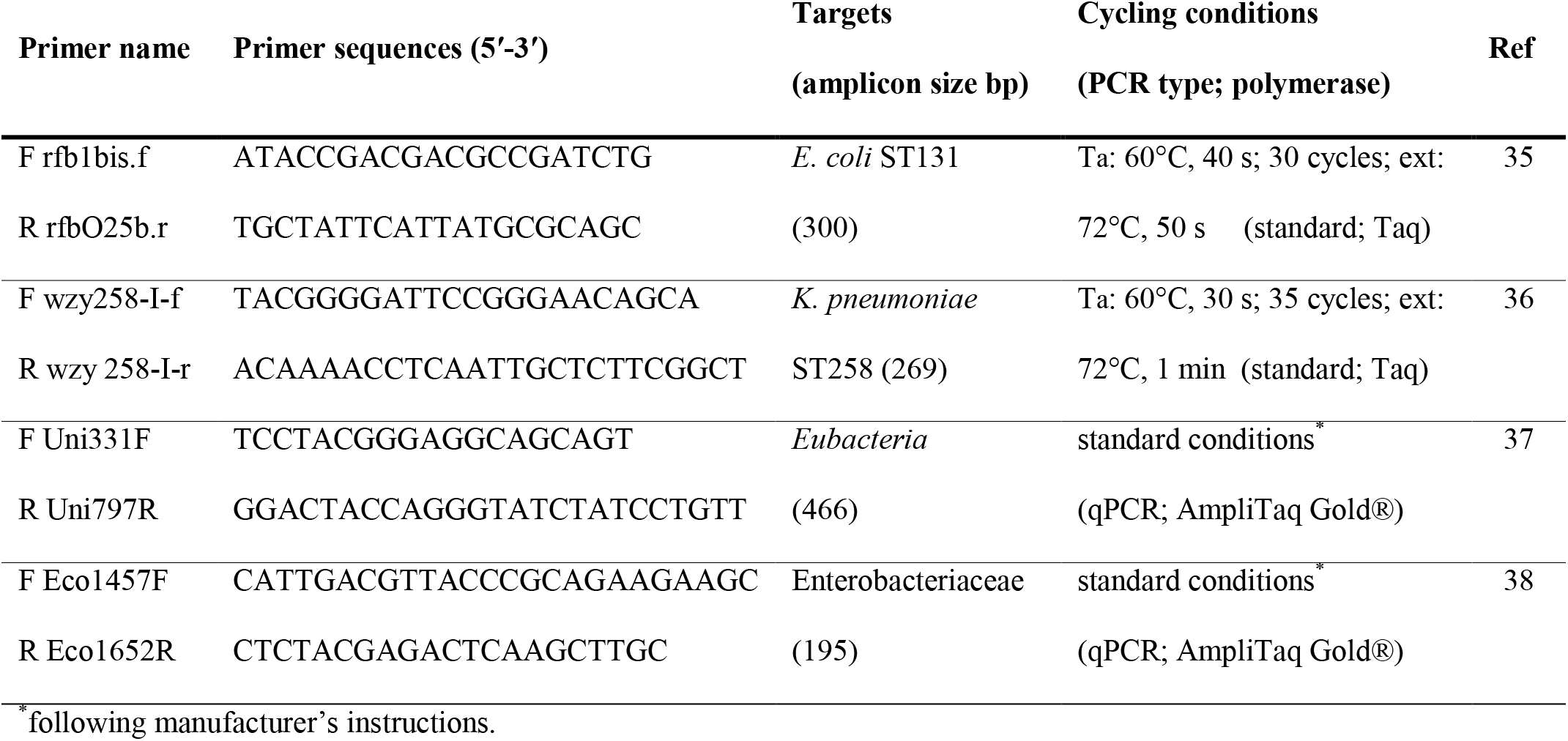
PCR conditions and primer sequences used in this study.

### Metagenomics

For 16S r-RNA gene amplicon sequencing, total microbial DNA was extracted from homogenized faecal pellets using the QIAamp Fast Stool Mini kit (Qiagen, Hilden, Germany) following manufacturer’s instructions. Sequencing was performed at the Ramaciotti Centre for Genomics (UNSW, Sydney, NSW, Australia) on an Illumina MiSeq platform (2 x 250 bp paired end reads) using the 515F and 806R primers for amplification of the V4 region of the 16S r-RNA encoding gene.^39^ Sequence clustering into OTUs (Operational Taxonomic Units) was performed using the QIIME 1.9.1 pipeline^40^ for closed reference OTU picking against the Greengenes database (97% sequence similarity)^41^ using the UCLUST algorithm.^42^ This study focussed on macro changes in gut microbial composition with analysis of most abundant OTUs. Therefore, the data was filtered to exclude singletons and rare OTUs and to include main phyla only (minimum total reads >8 M). Classified OTUs were used to calculate the relative abundance of bacterial groups (Phylum, Order, Family) in each sample. Diversity within samples as species richness (alpha diversity) was calculated using Shannon’s index of diversity at multiple rarefaction depths to ensure equality of sequence number for each sample and provide robustness to the analysis. Alpha diversity data presented here are for a rarefaction depth with a minimum of 2000 reads per sample. Principal coordinates analysis (PCoA) was used to assess community similarity among samples (beta diversity) using Unifrac distance metrics based on the generated phylogeny tree. Matrixes of beta-diversity were then visualized in a two-dimensional plot using Emperor in QIIME. Compositional changes in the microbiome were visualised using heatmaps and bar charts generated by the phyloseq package^43^ in R (version 3.5.1)^44^ and LEfSe.^45^

#### Sequence data accession number

All 16S r-RNA amplicon sequencing data and metadata are available through the Sequence Read Archive (SRA; NCBI) under accession no. PRJNA602745.

### qPCR assays

Real time qPCR (SYBR®Green Applied Biosystems™, CA, USA; RotorGene, Qiagen, Hilden, Germany) was performed on microbial DNA from day 0 (pre-treatment), day 5 and 9 (antibiotic), day 18 and 26 (recovery), and day 32 (endpoint) to measure the relative changes in total Enterobacteriaceae, as a calculated percentage of the average total Eubacteria. Primers used for qPCR are listed in Table 1. Final assay volumes of 12.5 μl were dispensed in triplicate into 96-well plates and each experiment repeated thrice. *E. coli* DNA of known concentration was used for standard curves and DNA from *Enterococcus ssp* (Gram-positive) and *Pseudomonas aeruginosa* (Gram-negative, Pseudomonadaceae) as controls.

### Statistical analysis

The mean bacterial load (CFU/g) for each group of mice (n=3) was log_10_ transformed and analyzed using PRISM, version 7 (GraphPad, San Diego, CA). Statistical analyses were performed in GenStat (18^th^ ed., VSN International, Hemstead, UK). The percentage of Enterobacteriaceae was log_10_ transformed for normality. One-way ANOVA at each time point was used with Fisher’s protected least significance difference test for multiple comparisons.

## Results

### Effects on microbiome composition vary with antibiotic treatment

All murine microbiomes pre-antibiotic treatment (pre-Ab, day 0) showed comparable composition at both the Phylum (Figure 1) and Order level (Figure 2), and no significant differences in OTU abundance were detected using LEftSe (P>0.05). Untreated (none) and sham (saline) treated mice presented comparable microbiome profiles at all stages (Figure 2) with predominance of Bacteriodetes and other obligate anaerobes (Clostridiales and Deferribacteriales), undisturbed during the study period (Figure 1 and 2).

**Figure 1.**
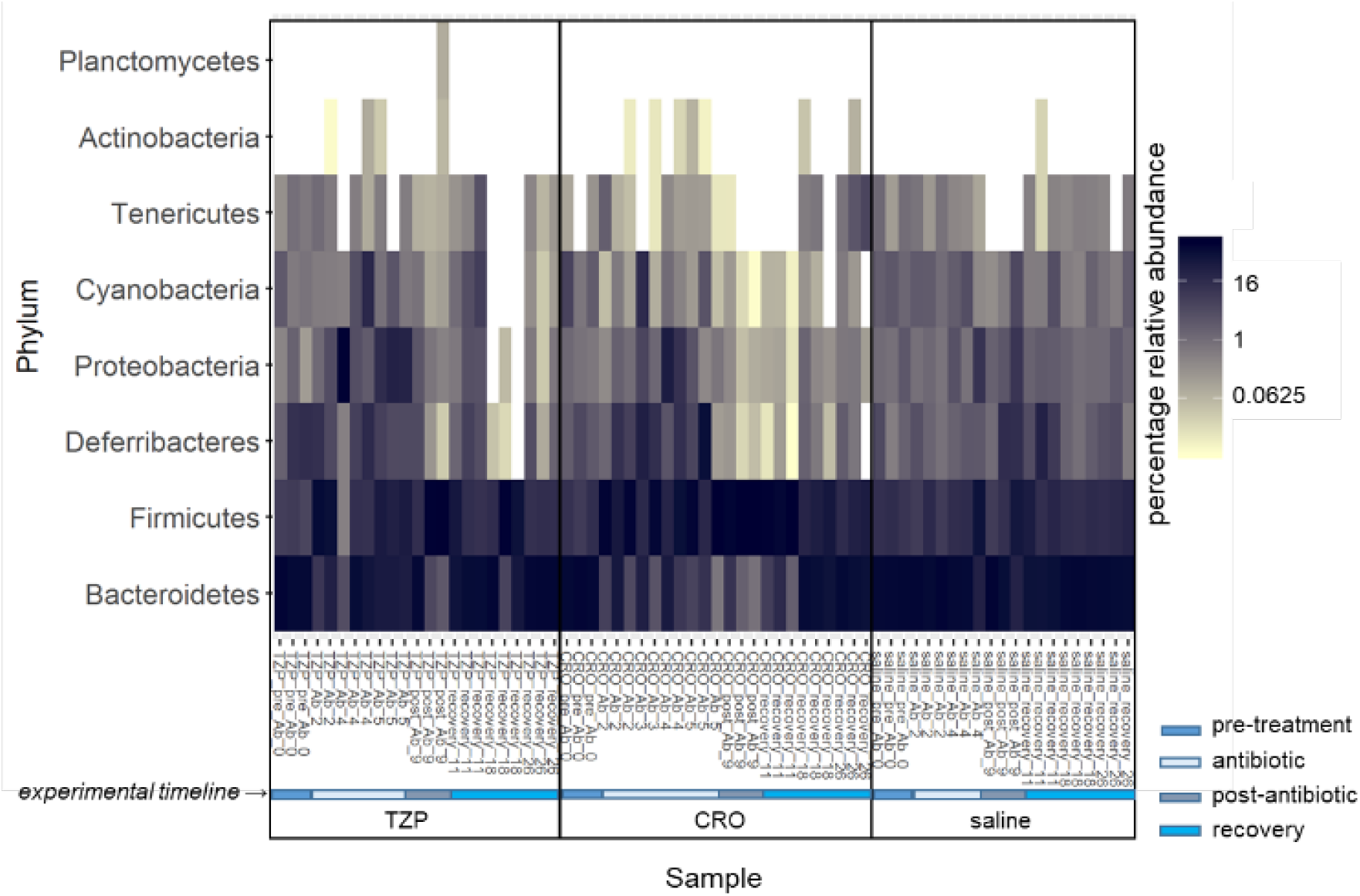
Microbiome composition (main phyla) of murine guts with and without antibiotic treatment. Compositional changes in Balb-C mice (n=3 per group) treated with antibiotics (TZP, piperacillin-tazobactam, or CRO, ceftriaxone) or sham treated (saline only) were compared by measuring the relative abundance of main phyla by 16S-rRNA gene analysis using QIIME-1.^40^ CRO-treated microbiomes showed the greatest variation post-antibiotic, with loss of Bacteroidetes and increase in Firmicutes and slower recovery to pre-antibiotic profiles. Total microbial component from faecal pellets was considered a proxy for the gut microbiome. Samples were collected prior to treatment at day 0 (pre_Ab_0), on days 2, 4 and 5 during treatment (Ab_2, Ab_4, Ab_5), post treatment on day 9 (post_Ab_9), and at various stages of recovery on days 11, 18 and 26 (recovery_11, recovery_18, recovery_26).

**Figure 2.**
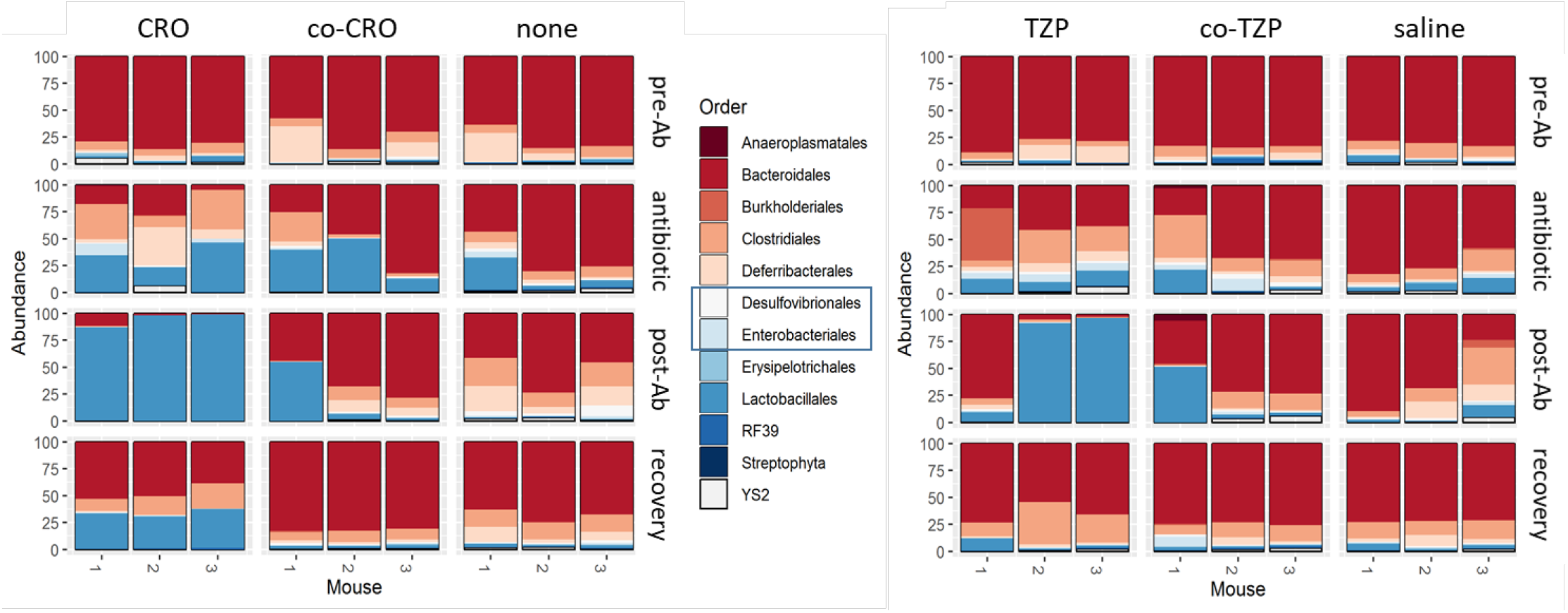
Changes in murine gut microbiome composition at the Order level after antibiotic treatment. Balb-C mice (n=3) were treated with either a penicillin-like antibiotic (TZP) or a third-generation cephalosporin (CRO) for 5 days and recovery monitored after cessation of antibiotic. Sham (saline) and no treatment (none) control groups received no antibiotics. One antibiotic-treated mouse (co-CRO-1 and co-TZP-1) was also co-housed with two untreated mice (co-CRO-2,3 and co-TZP-2,3). Relative abundance (%) at the Order level (pooled data for each timeframe: day 0 = pre-Ab; day 2, 4 and 5 = antibiotic; day 9 = post-Ab; day 11, 18 and 26 = recovery) was determined by analysis of 16S r-RNA amplicon sequencing (V4 region) data using QIIME-1.^40^

At the Phylum level, the microbiome composition of guts treated with TZP and CRO presented patterns unique to each antibiotic (Figure 1), but with common increase in the proportion of Lactobacillales (Firmicutes) and Proteobacteria during and post antibiotic administration and concomitant decrease of Bacteroidetes (Figure 1 and 2). In CRO-treated mice, this dysbiosis was sustained and murine microbiomes did not return to homeostasis (Figure 2), with Proteobacteria still significantly overrepresented at three weeks post treatment (recovery) (Figure S3). Treatment with TZP caused shifts in microbiome composition akin to those observed in CRO treated microbiomes with decrease in the levels of strictly anaerobic groups (Figure 2). However, TZP initially resulted in a more marked increase in Enterobacteriales than did CRO, but TZP treated microbiomes largely recovered to pre-antibiotic steady state within three weeks (Figure 1 and 2). Shifts in the relative abundance of Proteobacteria were detected in all groups only during or just after antibiotic treatment (Figure 2), and in antibiotic treated mice these were associated with increased diversification at family level (Figure 3). In CRO treated mice, the Alcaligenaceae family (Betaproteobacteria) was the predominant representative of the Proteobacteria during recovery (p<0.001) (Figure 3).

**Figure 3.**
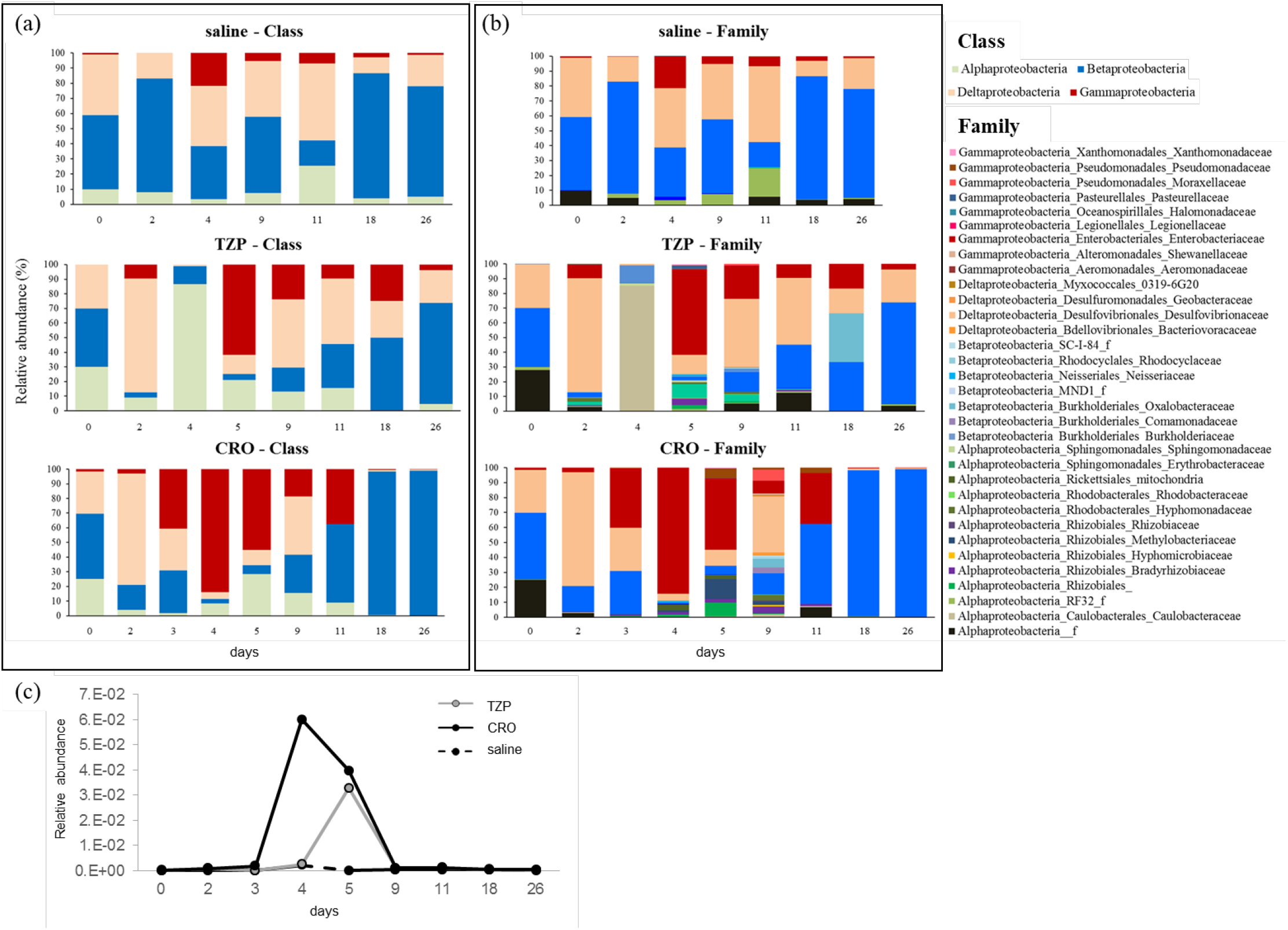
Proteobacteria relative abundance in murine microbiomes with and without antibiotic treatment. Microbiome composition at the (a) Class level, and (b) Family level, calculated as relative abundance of operational taxonomic units (OTU) from 16S-rRNA gene sequencing data. (c) Enterobacteriaceae relative abundance based on OTU counts in sequenced faecal DNA extracts. Mice were either treated with a penicillin-like antibiotic (TZP) or a third-generation cephalosporin (CRO) for 5 days and recovery monitored after cessation of antibiotic, or received a sham treatment but no antibiotic (saline).

Co-housing of an antibiotic treated mouse (co-CRO and co-TZP, mouse 1) with two untreated individuals (co-CRO and co-TZP, mice 2 and 3) promoted rapid recovery from dysbiosis (Figure 2). The microbiomes of co-housed untreated individuals (mice 2 and 3) resembled those of sham treated mice, while the microbiomes of treated co-housed mice (co-CRO and co-TZP mouse 1) were comparable to those of mice treated with the same antibiotic during and post treatment, returning to pre-treatment conditions within three weeks (Figure 2). Notably, a temporary dysbiosis was observed in the untreated mice co-housed with a CRO treated individual (co-CRO), but not in the co-TZP group (Figure 2).

### Enterobacteriaceae transiently increase in antibiotic treated murine microbiomes

Metagenomic analysis showed a 1-2 log increase in Enterobacteriaceae during (day 4/5; TZP P=0.003; CRO p<0.001) or immediately after antibiotics (day 9; CRO P=0.052) in all treated mice (Figure 3). A small increase was also observed in control mice (saline) on day 4 compared to day 2 levels (P=0.049) (Figure 3 and Figure S4). Prior to antibiotics, total Enterobacteria counts were comparable in all mice (~10^7^−10^8^ CFU/g), with levels remaining unchanged in controls, but decreasing below detection level in antibiotic treated mice (TZP and CRO) before recovery to pre-antibiotic values (~10^7^ CFU/g three weeks post antibiotic). In co-habiting treated mice (m1_co-TZP and m1_co-CRO), this increase was less pronounced than in other treated mice (Figure S4b). Levels of Enterobacteriaceae in co-habiting untreated mice also increased during antibiotic exposure and persisted in recovery in co-CRO mice (Figure S4b).

### Ceftriaxone caused a significant reduction in microbial diversity

A decrease in diversity in terms of species richness was observed following antibiotic treatment (Figure 4a). There was a significant (p<0.001) and protracted effect in the CRO group with limited recovery (day 11 to 26) when compared to sham and TZP treated microbiomes. A decrease in overall diversity was observed during TZP treatment and immediately afterwards (post_Ab, day 9; p<0.001), but levels during recovery (day 11 to 26) were comparable (p<0.001) to those in untreated individuals (Figure 4a). Beta-diversity indicators showed clustering of all pre-treatment samples with sham (saline) and no treatment (none) samples (Figure 4b). During antibiotic and immediately post treatment (post-Ab) CRO samples clustered with TZP samples separately from those of untreated mice. In agreement with compositional data, the CRO recovery samples partly grouped with other CRO samples and partly with saline, indicating only partial recovery to pre-antibiotic status. TZP recovery samples clustered following an independent trajectory towards the saline group (Figure 4b).

**Figure 4.**
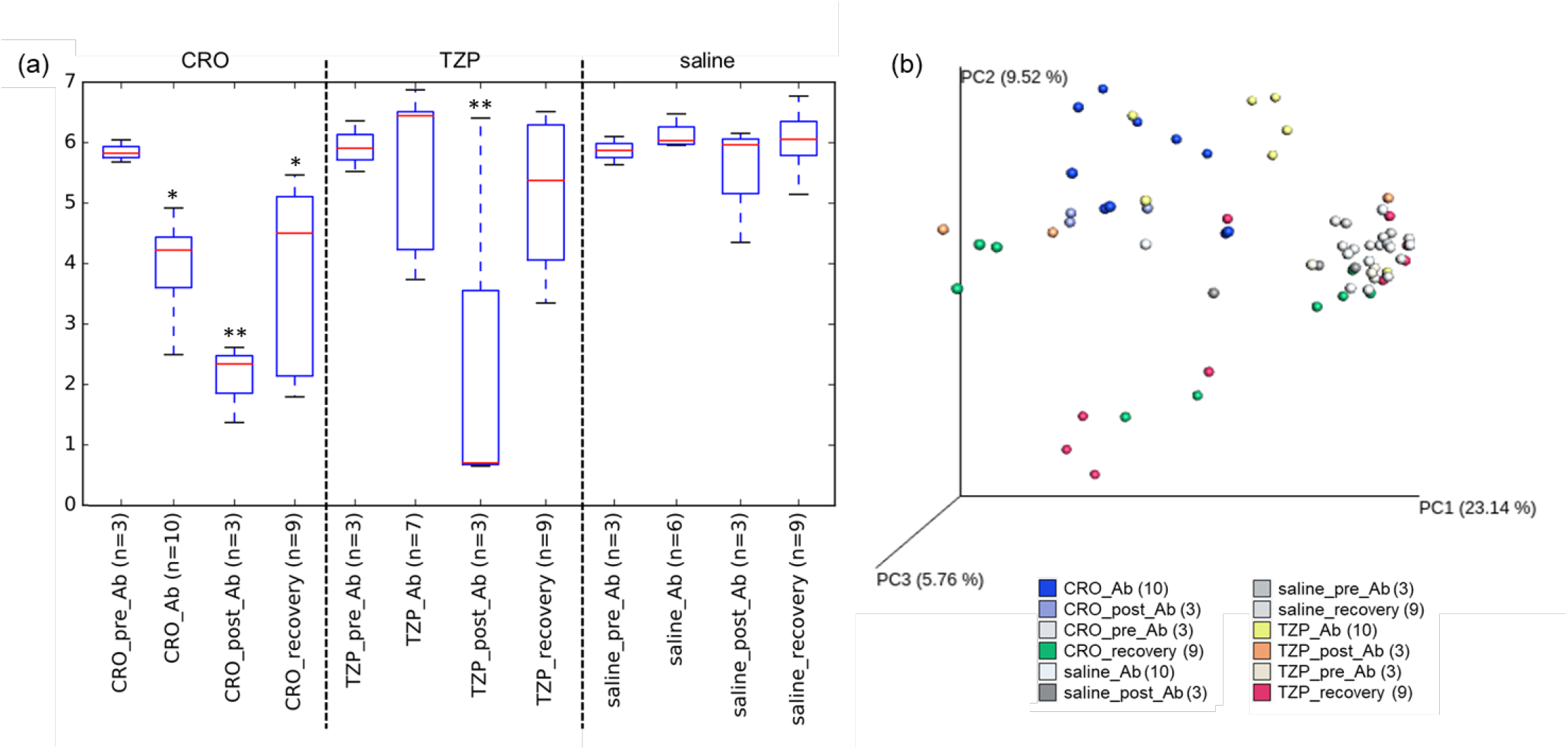
Diversity within and between murine microbiomes after antibiotic treatment. (a) Diversity in terms of species richness and evenness (alpha diversity) in faecal samples from mice treated with antibiotics (TZP, piperacillin-tazobactam; CRO, ceftriaxone) or sham treated (saline) was measured using the Shannon index. Shifts towards lower diversity can be observed during and after treatment with both antibiotics. *,** indicate significant difference (p<0.001). (b) Diversity between murine microbiomes (β-diversity) was assessed by principal component analysis (PCoA; Unweighted Unifrac distances) of microbial 16S-rRNA gene sequences in faecal samples, Pre-antibiotic samples clustered with no antibiotic (saline) samples, while samples from CRO treated and TZP treated mice (day 5, i.e. four days post-treatment; _Ab) clustered independently but on a similar trajectory. However, there was little indication of return to pre-antibiotic profiles in CRO-treated microbiomes (CRO_recovery; recovery week 3) when compared to TZP. Colored dots indicate composites of the microbial community during antibiotic (_Ab, day 3-5), immediately after antibiotic (_post_Ab, day 9), before antibiotic (_pre_Ab, day 0), and in recovery (1-4 weeks after antibiotic) (_recovery, days 11,18,26). Numbers in brackets indicate number of samples per group.

### Highly dysbiotic murine guts have reduced resistance to colonization with MDR *E. coli* and *K. pneumoniae*

Prior to bacterial inoculation (day 9), no CTX-resistant Gram-negative bacilli were detected on selective media used for detection of introduced *E. coli* ST131 and *K. pneumoniae* ST258 (MacConkey CTX8; limit of detection 2-2.5 log_10_ CFU/g) (Figure 5), though the presence of other ESBL Gram-negative Enterobacteria was observed on ChromAgar Van20-CTX8. Once colonization of dysbiotic guts was established (day 10), *E. coli* ST131 and *K. pneumoniae* ST258 persisted in all mice in significant amounts up to at least five days (day 14) post-inoculation (Figure 5). At two weeks post-inoculation (day 23), *E. coli* ST131 was only detected in one TZP-treated and one CRO-treated mouse (~10^6^ CFU/g), whereas *K. pneumoniae* ST258 was still detected in 2/3 TZP treated (~10^6^ and 10^3^ CFU/g) and all CRO-treated mice (~10^5^ CFU/g). At four weeks post-inoculation (day 38), levels of both pathogens dramatically decreased (~10^3^ CFU/g) (Figure 5). Colony PCR confirmed that all recovered CTX-resistant *E. coli* were ST131 and that CTX-resistant *K. pneumoniae* were ST258. At five weeks (day 44), no pathogens were detected. MDR *E. coli* and *K. pneumoniae* colonized untreated (no antibiotic; control) mice only transiently with no bacteria of either species detected by five days post inoculation (day 14).

**Figure 5.**
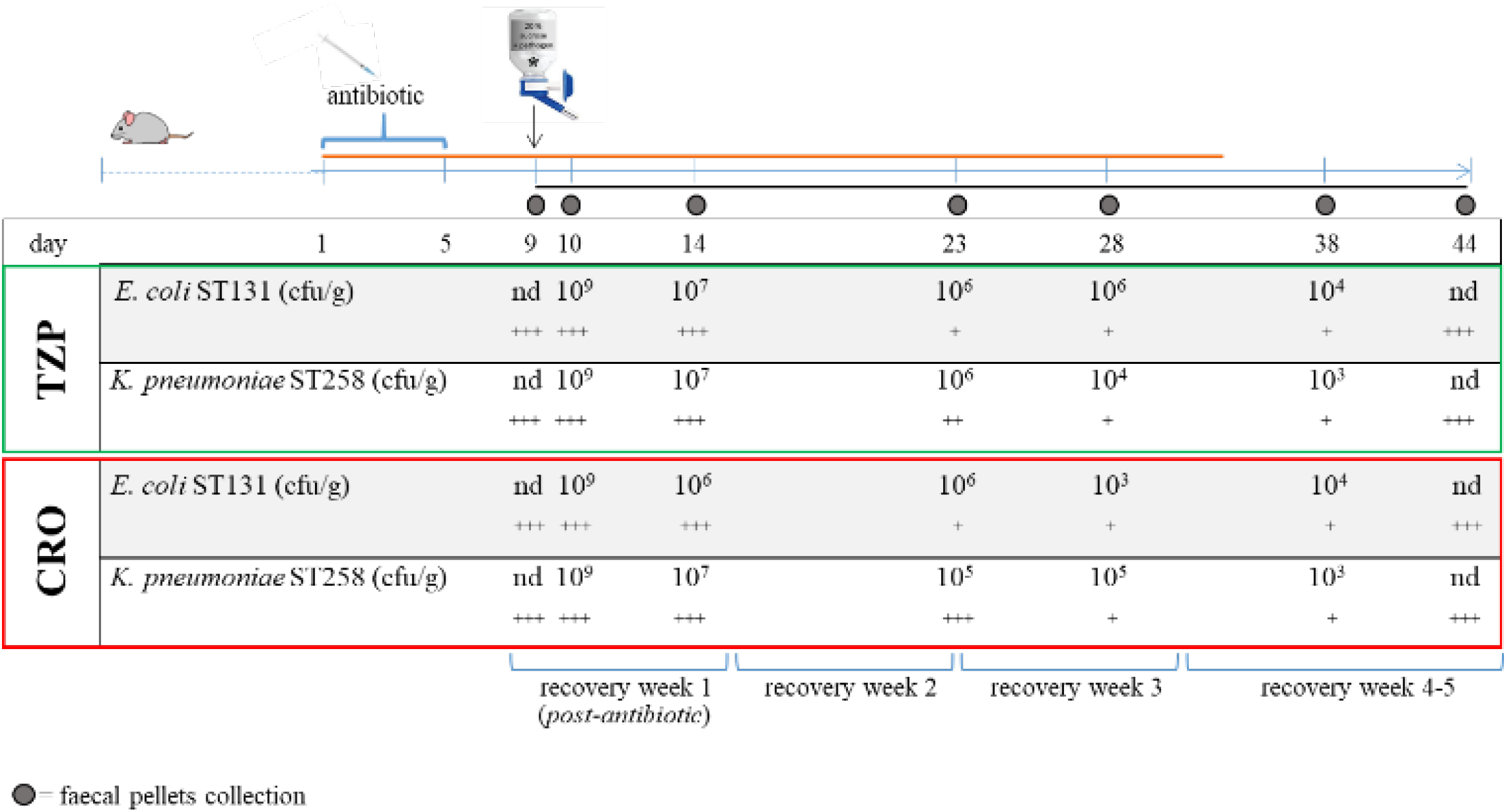
Colonization of murine guts treated with antibiotics with enteric opportunistic pathogens *E. coli* ST131 and *K. pneumoniae* ST258. nd, none detected; +, detected in 1/3 mice; ++, in 2/3 mice; +++, in 3/3 mice. Black line, “colonization resistance” experimental timeline (day 9 to 44). Orange line, “antibiotic effects” experiment timeline, ending at day 32. *pathogen colonization in drinking sucrose solution. TZP, piperacillin-tazobactam; CRO, ceftriaxone.

### Residual antibiotic activity in faeces was negligible

Prior to colonization, antibiotic activity was detected in all treated mice at 24 h post-antibiotic injection at approximately 8-32 μg/ml TZP and 0.5-1 μg/ml CRO (Figure S5). Antimicrobial activity was greater at 3 h than at 24 h after treatment with both antibiotics (Figure S5b and c). There was no indication of antimicrobial activity in our assay in faecal pellets collected three days after TZP treatment (day 8, Figure S5b). However, residual antimicrobial activity persisted for three days following CRO treatment in 3/6 mice and for four days in 2/6 mice (Figure S5b and c). On the day of colonization with invasive pathogens (day 10, five days post antibiotic treatment) there was no detectable activity of either antibiotic (Figure S5).

## Discussion

The effects of antibiotic treatment vary both in terms of antibiotic resistance transfer and collateral damage to bystander gut microflora depending on the drug used.^14,38,46^ Some antibiotics (e.g. ampicillin, ciprofloxacin, β-lactams, ceftriaxone) have been shown to impact gut microbiome composition with disruption of homeostasis associated with decreased resistance to pathogen colonization (infection)^8,12–15,46^ and compromised long-term wellbeing (immune system function).^47,48^ Two broad spectrum antibiotics, piperacillin/tazobactam (TZP) and ceftriaxone (CRO), have broadly comparable activity against medically important pathogens, and are routinely used in hospital inpatients, particularly in the critically ill.^5,16,24^ As only few studies had done before,^7,12,49^ we directly compared the effects of these two antibiotics, administered in doses equivalent to human therapy, on the composition of the murine gut microbiome to determine whether one antibiotic may provide a better clinical choice than the other.

With both TZP and CRO, we observed an expected relative shift in the proportion of the two phyla dominating the gut microbiome, Bacteriodetes and Firmicutes.^50^ The baseline composition of the murine gut microbiome depends mainly on diet^51^ and in our experiment the untreated gut flora of all mice was dominated by the Bacteriodetes, with both antibiotics promoting an overall decrease in the order Bacteroidales (mostly obligate anaerobes) and a converse increase in Lactobacillales and/or Clostridiales (phylum Firmicutes). However, most importantly, specific compositional shifts were unique to each antibiotic.^52,53^ CRO had a more pronounced negative impact on the diversity of the gut microbial community than TZP with a significant and prolonged decrease in both species richness and abundance, and slower recovery towards steady state.

Proteobacteria overall increased with both antibiotics, mainly during treatment, i.e. under direct antibiotic selective pressure and at the height of dysbiosis (days 4 & 5), though levels of the Enterobacteriaceae (Gammaproteobacteria), the family most commonly chosen as dysbiosis indicator, only rose transiently during, or immediately after, treatment with both drugs. Antibiotic-associated dysbiosis raises levels of available oxygen in the gut lumen, favouring amplification of facultative aerobes, such as the Enterobacteriaceae and members of the Betaproteobacteria.^10–14,50,53,54^ Previously, significant blooms in the Enterobacteriaceae were observed following the use of antibiotic cocktails, oral administration protocols or sub-inhibitory dosing of various single antibiotics.^7,21,50,52–55^ Our study endeavoured to use a mouse model where drug administration more closely resembled clinical human protocols with comparable dosing. Though TZP was administered in single dose, instead of multiple as customary in clinical practice, which could mask some initial dysbiotic signals, there was no evidence of residual antibiotic activity past the end of antibiotic treatment that could have led to bias. Route and duration of antibiotic administration as well as the strategies used for detection (16S-rRNA amplicon etc.) and sampling (single or multiple time point etc.) are known to influence observed microbial shifts and may account for the discrepancies between studies. We showed that TZP and CRO stimulate initial diversification of Proteobacteria and our findings suggest that levels of total Proteobacteria may be a more reliable marker of the differential impact of antibiotics than those of the Enterobacteriaceae alone. Studies that are specifically interested in investigating Enterobacteriaceae fluctuations in humans, may therefore benefit from animal models that more closely reproduce clinical treatment protocols like ours.^21,53^

In this study, we also tested susceptibility to invasion by opportunistic MDR pathogens at the height of antibiotic-induced dysbiosis, showing that an imbalanced gut microflora is prone to prolonged colonization independently from the specific antibiotic used. MDR *E. coli* ST131 and *K. pneumoniae* ST258, which are apt colonizers of the healthy human gut, have been identified as major agents of both community acquired and nosocomial infections.^56^ Establishment of these key pathogens in murine guts has been linked to antibiotic-induced decrease in anaerobic flora.^14,21–23,50–53,57^ ESBL *K. pneumoniae* colonization has been associated with piperacillin-tazobactam, ceftriaxone, and ceftazidime when administered in large doses.^29^ We know that the invading *E. coli* and *K. pneumoniae* strains used in our study do not persist in healthy murine guts without antibiotic pressure (controls here and ^34^), indicating that their persistence was a direct consequence of dysbiosis. Loss of colonization resistance is pathogen and disturbance dependent,^28,58^ with different bacterial strains having variable propensity for persistent colonization in a context-dependent manner based on the community structure after depletion of specific beneficial microbes by specific antibiotics. The inhibitory activity of TZP against ESBL-producing organisms (e.g. *E. coli* ST131) may be sufficient to prevent initial establishment of colonization, but, as shown here, in guts with disrupted homeostasis (e.g. ICU patients) where competition from indigenous flora is reduced, the use of this antibiotic may still lead to persistent high-density colonization by exogenous clones and potential contribution to antibiotic resistance transmission.

Our work shows that CRO use promotes a long-term dysbiosis with expansion and diversification of Proteobacteria, as well as reduced microbial diversity and slower recovery of murine microbiomes to pre-treatment conditions, when compared to TZP, indicating that the use of penicillin-like antibiotics instead of CRO should be recommended where possible, even though both antibiotics may have similar detrimental effects on colonization resistance.

## Acknowledgements

We acknowledge the Ramaciotti Centre for Genomics (RCG) at the University of New South Wales Sydney as the service provider for our microbiome sequencing data. We thank Dr Nouri L. Ben Zakour for her bioinformatics support.

## Funding

This work was supported by the Australian National Health and Medical Research Council (NHMRC) (GRP1046889) to JI.

## Transparency Declaration

None to declare.

## Notes

### Competing Interest Statement

The authors have declared no competing interest.

